# Discrete Protein Metric (DPM): A new image similarity metric to calculate accuracy of deep learning-generated cell focal adhesion predictions

**DOI:** 10.1101/2021.12.10.472147

**Authors:** Miguel Contreras, William Bachman, David S. Long

## Abstract

Understanding cell behaviors can provide new knowledge on the development of different pathologies. Focal adhesion (FA) sites are important sub-cellular structures that are involved in these processes. To better facilitate the study of FA sites, deep learning (DL) can be used to predict FA site morphology based on limited datasets (*e*.*g*., cell membrane images). However, calculating the accuracy score of these predictions can be challenging due to the discrete/point pattern like nature of FA sites. In the present work, a new image similarity metric, discrete protein metric (DPM), was developed to calculate FA prediction accuracy. This metric measures differences in distribution (*d*), shape/size (*s*), and angle (*a*) of FA sites between the predicted image and its ground truth image. Performance of the DPM was evaluated by comparing it to three other commonly used image similarity metrics: Pearson correlation coefficient (PCC), feature similarity index (FSIM), and Intersection over Union (IoU). A sensitivity analysis was performed by comparing changes in each metric value due to quantifiable changes in FA site location, number, aspect ratio, area, or orientation. Furthermore, accuracy score of DL-generated predictions was calculated using all four metrics to compare their ability to capture variation across samples. Results showed better sensitivity and range of variation for DPM compared to the other metrics tested. Most importantly, DPM had the ability to determine which FA predictions were quantitatively more accurate and consistent with qualitative assessments. The proposed DPM hence provides a method to validate DL-generated FA predictions and can be extended to evaluating other predicted or segmented discrete structures of biomedical relevance.

## Introduction

To better understand human health, gaining insight into different cell behaviors is key. Cells adapt to new situations by perceiving information and transforming it into changes in their gene and protein expression repertoire (Tosh & Slack, 2002). These changes result in responses that could lead to different pathologies. One such example is the way cells sense and respond to their mechanical environment. Mechanosensing, how a cell senses and responds to inter- and intra-cellular stresses, can provide new knowledge on the development of diseases such as cardiovascular disease (Gaetani et al., 2020), cancer progression (Kärki & Tojkander, 2021), or fibrosis (Tschumperlin et al., 2018). These behaviors may be explained in part through close observation of cell morphology (McCarron et al., 2017). In response to stimuli (*e*.*g*., change in substrate stiffness, fluid shear stress), a single cell or group of cells make dynamic morphological adjustments. These morphological changes are accompanied with changes in cell adhesion to the extracellular matrix and force distributions throughout the cell (Yeung et al., 2005). Insight into both changes can be provided by the properties of a sub-cellular structure known as focal adhesions (FA). FA sites consist of large macromolecular assemblies of proteins that associate with integrin to provide anchor points for a cell to adhere to the extracellular matrix. FA sites play a fundamental role in force sensing, which influences several cellular processes and functions, including cell migration and cell cycle (Haase et al., 2014). Therefore, studying the morphology of these FA sites can provide a better understanding of cell behavior. Such morphology can show changes in FA site number, size, and location over time (Berginski et al., 2011) or can show changes in FA site alignment due to external stimuli such as shear stress (Davies et al., 1994).

One method that has been used to quantify the change to sub-cellular morphological features (*i*.*e*., shape, size, intensity, and texture) is image-based profiling. In this method, averaged profiling is used to summarize a cell population (a sample) into a fixed length vector (a sample’s profile), with one value per feature per sample (Rohban et al., 2019). Image-based profiling has been used by studies coupling morphological features with gene expression, where the modeling procedure involves the selection of genes that are associated with a given image-based feature (Nassiri & McCall, 2018). However using this method, multiple configurations of distinct subpopulations of cells could yield identical average profiles (Rohban et al., 2019). As a result, this method is limited in its ability to recognize cell heterogeneity. Therefore, new methods are needed to quantify morphology.

Machine learning (ML) is a computational method that has been used recently to analyze the morphology of cells. For example, deep learning (DL) algorithms, a subset of ML techniques, have been developed to perform cell segmentation in 2D microscopy images by detecting cells, separating touching cells, and segmenting sub-cellular compartments (*e*.*g*., nucleus and cytoplasm) (Al-Kofahi et al., 2018). ML has also been used to classify cell images according to their morphological features, where these features have been previously extracted from image segmentation (Rodellar et al., 2018). While these methods provide great insight into cell morphology, they have not been applied to study all sub-cellular components. For the study of cells, it is necessary to generate models that identify the morphology of the key components involved in their function, including the FA sites. A potential use of DL on morphological analysis is through *in silico* labeling of cellular components. This approach has been proved to be effective through the development of DL algorithms that predict fluorescence labels (*e*.*g*., nuclei and cell membrane) on unlabeled cell images (Christiansen et al., 2018) or DL networks that predict sub-cellular protein structure images (*e*.*g*., alpha actinin, alpha tubulin) using other cell label images (*e*.*g*., nucleus and cell membrane) as input (Yuan et al., 2019). These algorithms provide the advantage of being highly adaptable. For instance, through transfer learning the algorithms can use knowledge acquired from previous experiences to perform a new task. This approach is especially beneficial when there is limited amount of annotated data available (Kensert et al., 2019). The use of transfer learning on these types of algorithms has the potential to predict the morphology of relevant sub-cellular structures, such as the FA sites, using limited data (*e*.*g*., only cell membrane).

Although FA sites can be predicted from limited data, validating DL-generated predictions of FA sites can be a challenging task. Such validation would need to be carried out by measuring the similarity between the FA ground truth image of a cell and the FA image DL-predicted. Multiple image similarity metrics have been used for sub-cellular predictions. For example, Pearson correlation coefficient (PCC) has been used to measure nuclei fluorescence image prediction accuracy (Christiansen et al., 2018), feature similarity index (FSIM) has been used for delineation and detection of cell nuclei (John et al., 2016), and Intersection over Union (IoU) have been used to measure accuracy of semantic segmentation to identify nuclei in microscopy cell images (Punn & Agarwal, 2020). These metrics are better suited to compute the image similarity of large sub-cellular structures such as the nucleus. However, for FA predictions, accuracy scores can be difficult to compute and interpret given that FA sites are much smaller structures and have a discrete/point-pattern nature. FA predictions will be at different locations, differ in number, and will have different aspect ratios, areas, and angles of orientation, so a single metric might fail to simultaneously capture variations in all of these measurements. Furthermore, most existing metrics yield scores that are inconsistent with qualitative assessment. Therefore, the objective of this research is to develop a new image similarity metric to evaluate DL-generated FA sites predictions. The new metric, discrete protein metric (DPM), measures similarity in distribution (*d*), shape (*i*.*e*., aspect ratio) and shape/size (*i*.*e*., area) (*s*), and orientation angle (*a*) of FA sites. The DPM was compared to three commonly used image similarity metrics for cell images (PCC, FSIM, and IoU) and tested on DL-generated FA predictions for validation of its effectiveness on calculating FA predictions accuracy scores that are in better agreement to qualitative assessment.

## Methods

### 1. Cell culture and culture media

Human Microvascular Endothelial cells (HMEC-1, #CRL-3243, ATCC, Manassas, VA) were maintained in complete growth media. Complete growth media consisted of MCDB 131 media (#10372019, Gibco, Grand Island, NY) supplemented with 2 mM L-glutamine (#25030081, Thermo Fisher Scientific), 10% (v/v) FBS (#15000044, Thermo Fisher Scientific) and penicillin/streptomycin (100 U/ml and 100 μg/ml concentration, respectively). Cells were seeded at a density of 10,000 cells/cm^2^ onto fibronectin-coated glass coverslips (#CS-25R17, thickness 1.5, Warner Instruments, Hamden CT) (fibronectin, 20 μg/ml, #33016-015, Thermo Fisher Scientific) mounted in 6-well plates. The cells were grown to confluence in a humidified environment at 37°C with 5% CO_2_.

### 2. Immunofluorescence labeling

Once the cells reached confluence, the plasma membrane of live cells were stained with Wheat Germ Agglutinin (WGA) (CF633, Biotium, Fremont, CA). Cells were washed twice in Hank’s balanced salt solution (HBSS, #14025076, Thermo Fisher Scientific), then incubated with WGA (5 μg/ml) for 30 minutes at 37°C, then washed twice in HBSS. Next, the cells were simultaneously fixed (4% (w/v) paraformaldehyde (PFA) in PBS) and permeabilized in Triton X-100 (0.5% (v/v), 5 min, #T9284, Sigma-Aldrich). Then post-fixed in 4% (w/v) PFA in PBS, followed by PBS wash (3 × 5 min). To image the nucleus, cells were stained with Hoecsht 33258 (1:1000, #116M4139V, Sigma-Aldrich). A blocking solution comprised of goat serum (1:20, #G9023, Sigma-Aldrich), 0.1% Triton X-100, and 0.3M glycine in 0.1% (w/v) BSA was added to cells for 30 minutes. To image FA sites, cells were blocked with the previously mentioned blocking solution for 30 minutes at room temperature. Then cells were incubated overnight with anti-paxillin (1:500, Abcam, #ab32084). This was followed by a 2-hour incubation with secondary antibody goat anti-rabbit Alexa Fluor 488 (1:500, #ab150077, Abcam, Waltham, MA, USA), and a 0.1% (w/v) BSA wash (3 × 5 min). The fibronectin coated cover slips were then removed from the well plates and mounted cell-side-down onto individual glass microscope slides using ProLong Glass (#P36982, Thermo Fisher Scientific).

### 3. Microscopy and image acquisition

A Leica TCS SP5 confocal microscope with a 63×/1.40 NA oil immersion lens was used to image the monolayer. UV 405 nm (nucleus), UV 488 nm (FA sites), and a HeNe 633 nm (membrane) lasers were used to sequentially excite samples. Images were acquired at 2048 pixels × 2048 pixels, with an *x*-*y* spatial resolution of 0.132 μm/pixel and *z* spatial resolution within the range of 0.1-0.15 μm/slice for different imaging sessions. A total of 8 fields of view (*i*.*e*., 8 image stacks) were acquired from cells belonging to all well plates, from which the slice with the brightest membrane and FA signal was taken.

### 4. Image processing and data augmentation

For each field of view, an overlay of the red color channel for membrane and the green color channel for FA sites was obtained and processed as an RGB image. From each image, individual cells were manually segmented using *ImageJ ver1*.*52i* (Schneider et al., 2012). A total of 60 single-cell images were obtained from all 8 fields of view. Then, the dataset formed by these individual cells was augmented performing random rotation and translation for eight iterations. For each iteration, a random angle between 0° and 360° and a random vertical and horizontal shift between -100 pixels and 100 pixels were chosen to rotate and shift each individual cell. This resulted in a total of 480 single-cell images. Each image was rescaled to a size of 256 pixels × 256 pixels. Finally, FA color channel was processed using a top hat filter with a 3 × 3 rectangular kernel to eliminate background noise.

### 5. Data combination and transformation

The augmented single-cell images were combined with a dataset from the Allen Institute for Cell Science (Johnson et al., 2017) to increase the size of the dataset and use transfer learning for neural network training. This Allen Institute dataset consisted of 6077 single cell images containing cell membrane, nucleus, and one protein of interest stained. This dataset was divided into 10 classes, each class consisting of one protein of interest. One of these classes was randomly dropped and replaced with the acquired dataset. For all cells from the Allen Institute, the nucleus color channel was dropped, and each image was rescaled to a size of 256 pixels × 256 pixels. The final dataset consisted of 6077 images containing membrane and the protein/structure of interest. The images were then converted to HDF5 format. For the conversion, the images were transformed into NumPy arrays. The pixel values were normalized from the 0-255 range to a 0-1 range. The images containing both membrane and structure of interest were appended to a single array of dimensions (6077, 256, 256, 3) with the third color channel being empty. The labels (*i*.*e*., structure of interest) were transformed into a one-hot encoded vector and appended to a single array of dimensions (6077, 10). The color channel for cell membrane structure was separated from the images and appended to a single array of dimensions (6077, 256, 256, 2) with the second color channel being empty. Finally, the three arrays were stored into a single HDF5 file.

### 6. Neural network architecture and training

An open-source neural network (Yuan et al., 2019) was trained to predict the different structures of interest, using only cell membrane as input. For the objective of this study, only FA sites predictions were analyzed. The network was based on a conditional generative adversarial network (cGAN) architecture, consisting of a generator and discriminator (Yuan et al., 2019). The dataset was split into training and testing set using an 80/20 ratio. For the images of interest (*i*.*e*., FA sites) this resulted on 390 images for training and 90 for testing. The network was trained using the original parameters (Yuan et al., 2019) with a learning rate of 2 × 10^−4^ and a batch size of 10. The training lasted 50 epochs, as no improvements were observed after this point.

### 7. Discrete protein metric (DPM)

After training the network, information was extracted from predictions across the 90 samples in the test set, along with their respective ground truth. For such extraction, membrane and FA color channels from each prediction/ground truth pair were separated and binarized. For membrane images, the outline was segmented to produce an image of the segmented membrane. For FA images, FA sites that had an area < 1μm^2^ were dropped (Mullen et al., 2014) and the outline of each FA site remaining was segmented to produce an image of the segmented FAs. Subsequently, the *x-y* centroid coordinates for each FA site were extracted to describe their number and location and an image was produced. Furthermore, for each FA site a bounding box was drawn around them to describe their shape (*i*.*e*., aspect ratio) and size (*i*.*e*., area) and an image was produced. A schematic of this process can be seen in Fig. 1 and Fig. 2.

**Fig. 1:**
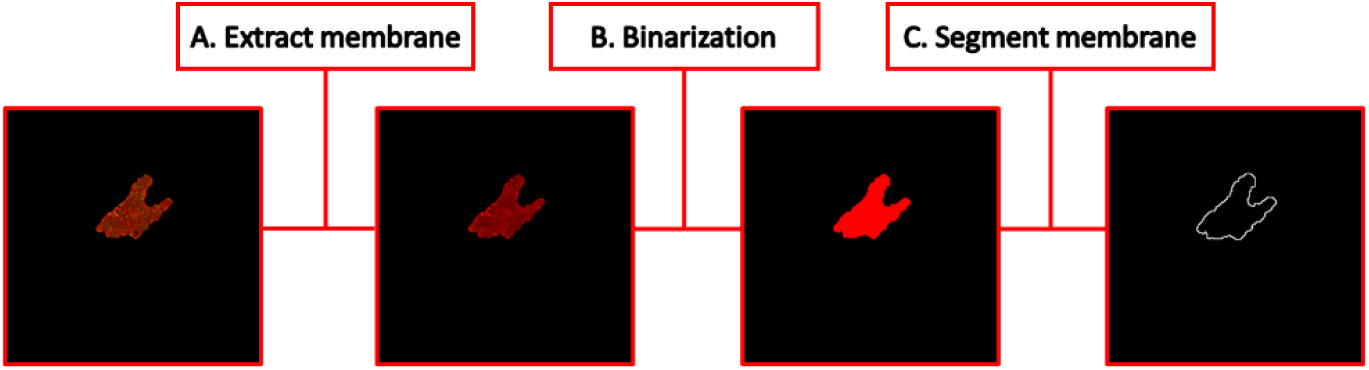
Information extraction pipeline of the the cell membrane segmentation process: (A) Membrane color channel extraction from cell image; (B) Binarization of membrane image; (C) Segmentation of membrane outline.

**Fig. 2:**
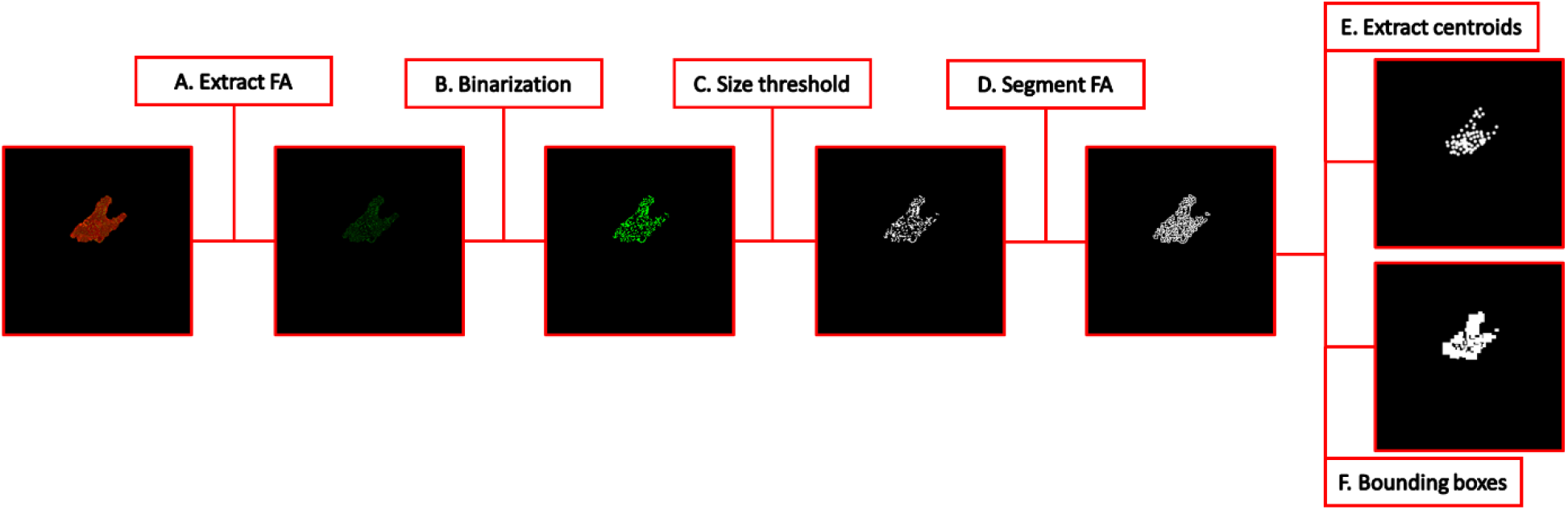
Information extraction pipeline of the FA site segmentation: (A) FA color channel extraction from cell image; (B) Binarization of FA image; (C) Size threshold where FAs with < 1 μm^2^ area were dropped; (D) Segmentation of FAs outline; (E) Extraction of centroids for each FA; (F) Bounding box drawn for each FA.

A new metric, discrete protein metric (DPM), was developed to use the information extracted to compute the accuracy of FA predictions. This metric accounts for three key aspects of a prediction: distribution (*d*), shape and size (*i*.*e*., aspect ratio and area) (*s*), and orientation angle (*a*) of FA sites. The calculation of DPM is a weighted sum of these three components. Equation (1) shows the calculation of the metric,

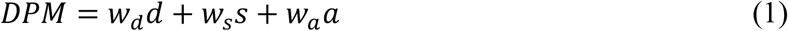

where *w*_*d*_, *w*_*s*_, and *w*_*a*_ are the relative weights for distribution, shape/size, and orientation angle respectively. These weights are assigned according to the relative importance of each component for the research question of interest. The sum of these weights must always equal one. For the purpose of later calculations, the importance of all measurements was considered to be equal (*i*.*e*., all weights were set equal to 0.33).

#### 7.1 Distribution (*d*)

For the distribution measurement, a *k*-means clustering algorithm (Pedregosa et al., 2011) was employed. The algorithm was trained individually for each ground truth FA site image in the test set. For such training, the coordinates of the centroids of each FA site were used as input to group them into five clusters. After training, the algorithm was used to predict which cluster each predicted FA site belonged to. If any predicted FA site was outside of the membrane outline, it was dropped. The number of FAs dropped was used as a penalizing factor to the overall *d* score. Finally, the ratio of number of FAs belonging to a particular cluster in the prediction to ground truth was calculated, and the average of ratios across all clusters was taken. Equation (2) shows the calculation of the distribution measurement,

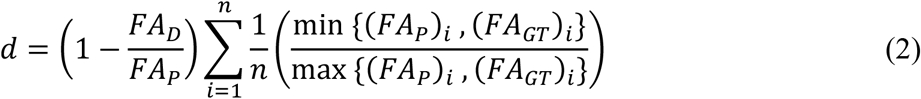

where *FA*_*D*_ is the number of FAs dropped, *FA*_*P*_ is the total number of FAs predicted, *n* is the number of clusters, and (*FA*_*P*_)_*i*_ and (*FA*_*GT*_)_*i*_ are the number of ground truth and predicted FAs in the *i*^th^ cluster respectively. For better interpretability of the *d* measurement, heat maps showing FA numbers and ratios within each cluster were generated. A schematic of the clusters formed after training the *k-*means algorithm can be seen in Fig. 3.

**Fig. 3:**
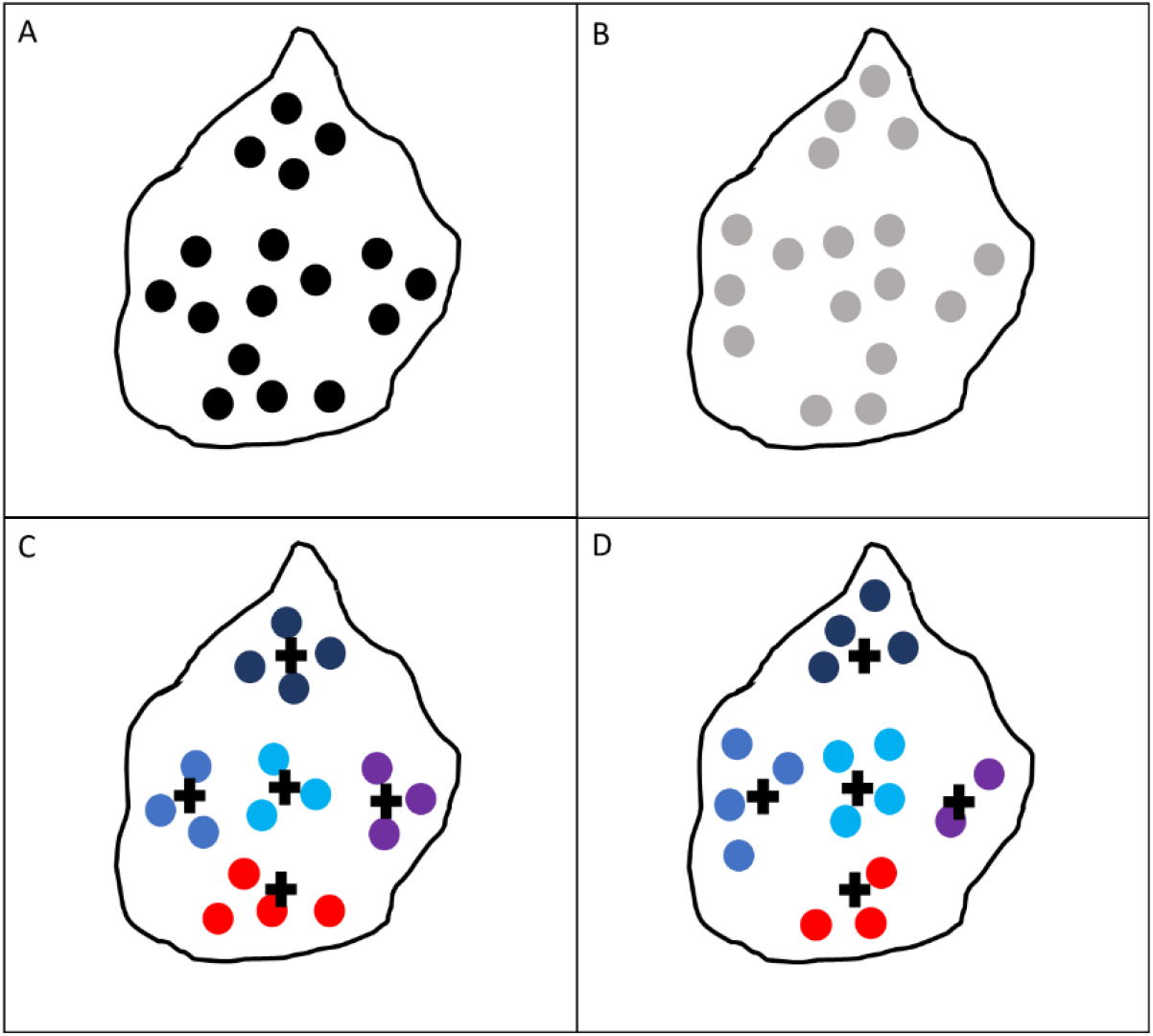
Schematic of clusters formed after *k-*means algorithm training: (A) Ground truth FA centroids; (B) Predicted FA centroids; (C) Clusters generated from ground truth data with each color representing a different cluster and cross marks representing the centroid of each cluster; (D) Predicted FA centroids assigned to their nearest cluster.

#### 7.2 Shape/size (*s*)

For the shape/size measurement, the location of the bounding box of each FA site in the prediction was matched to its nearest neighbor in the ground truth by setting their centroid coordinates equal to each other. After all boxes were at matching locations, their overlap was measured using an F1 score based on precision and recall. Equations (3), (4), and (5) show, respectively, the calculations of F1 score, precision, and recall,

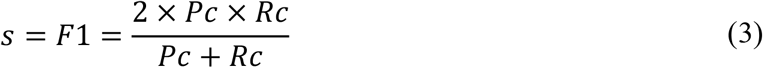

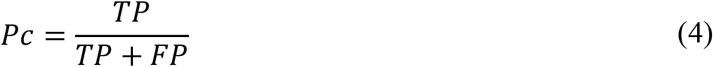

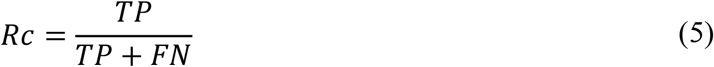

where *Pc* is precision, *Rc* is recall, *TP* is true positives, *FP* is false positives, and *FN* is false negatives. A schematic of the calculation of the *s* measurement can be seen in Fig. 4.

**Fig. 4:**
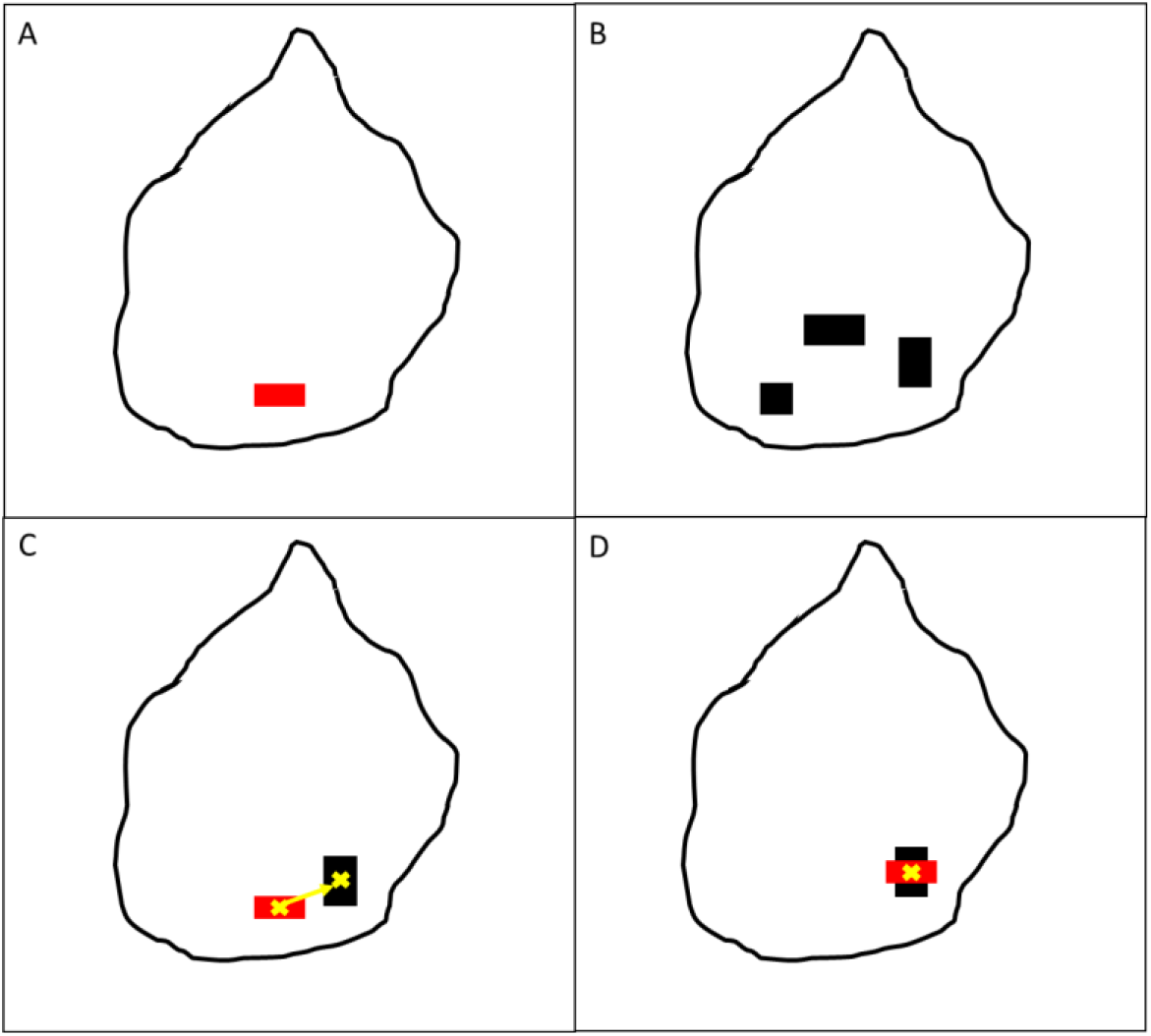
Schematic of calculation of *s* measurement: (A) Single prediction for a FA site bounding box represented in red; (B) Ground truth FA site bounding boxes on predictions vicinity in black; (C) Nearest neighbor identification with centroid for each FA represented by cross mark; (D) Location match using centroid coordinates.

#### 7.3 Orientation angle (*a*)

For the orientation angle measurement, the orientation angle of each FA site bounding box in the prediction was compared to the orientation angle of the nearest bounding box in the ground truth. To calculate such angles, the minimum area rectangle was calculated and the smallest angle that it formed with respect to the horizontal axis was taken. After angles were calculated, the deviation in the orientation angle was taken as the difference between the angle of the predicted box and the ground truth box. This difference was divided by 90° as this was considered to be the maximum difference there could be between the two angles. Equation (6) shows the calculation of the angle measurement,

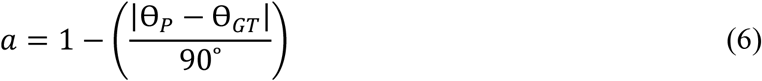

where θ_*P*_ and θ_*GT*_ are the rotation angles in degrees for prediction and ground truth, respectively. A schematic of the calculation of the *a* measurement can be seen in Fig. 5.

**Fig. 5:**
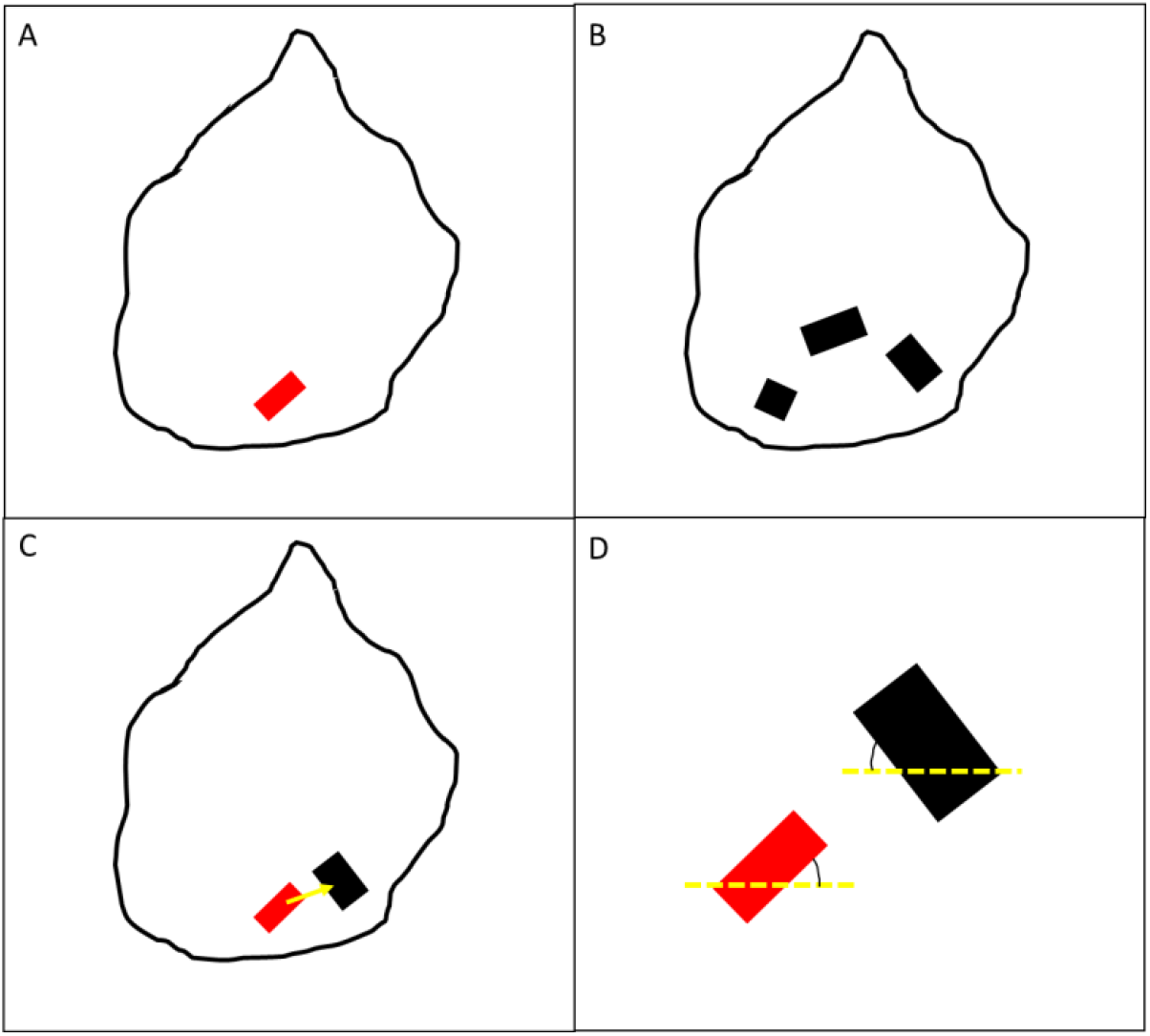
Schematic of the calculation of *a* measurement: (A) Single prediction for a FA site minimum area rectangle in red; (B) Ground truth FA site minimum area rectangles in the prediction vicinity in black; (C) Nearest neighbor identification; (D) Measurement of smallest orientation angle with respect to common horizontal axis.

### 8. Benchmark with image similarity metrics

To test the performance of the DPM, a benchmark with three common image similarity metrics was performed: Pearson correlation coefficient (PCC), Intersection over Union (IoU), and feature similarity index metric (FSIM). The metrics were compared by looking at the distribution of prediction accuracy across all 90 samples in the test set and by performing a sensitivity analysis.

#### 8.1 Distribution of prediction metrics and qualitative assessment

To look at the distribution of prediction accuracy, all four metrics were computed for each of the 90 predictions in the test set with respect to their respective ground truth image. The metric value for each cell was plotted, and the average was calculated as the intercept of a linear fit and the range of variation as the difference between minimum and maximum values. Furthermore, in order to determine if quantitave prediction accuracy was in agreement with quality of predictions, a qualitative assessment was performed. This assessment looked at differences between a ground truth image and its predicted image by identifying areas with different FA site densities (*i*.*e*., areas missing predicting FA sites or areas with overpredicted number of FA sites), with differences in FA size, and area with differences in FA site angles.

#### 8.2 Sensitivity analysis

A sensitivity analysis was performed by looking at the change in a metric value due to a quantifiable variation in either number, location, area, aspect ratio, or orientation of FAs predicted. Therefore, sensitivity (*S*) was defined as the difference in metric value for every 1% variation. Equation (7) shows the calculation of sensitivity,

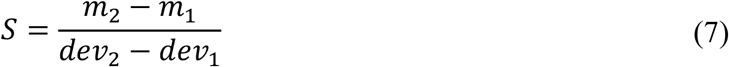

where *m*_2_ and *m*_1_ are the metric values at *dev*_2_ and *dev*_1_ variation values, respectively. For sensitivity of number of FAs, random FA sites were dropped from ground truth images in the test set at 5% intervals from 0-100%, where number of FAs dropped was treated as a percentage of the total number of FAs in a cell. Similarly, FAs were added at random locations inside the cell boundary and sensitivity to these changes was calculated. For sensitivity of location, centroid coordinates of FA sites in ground truth were randomly altered at intervals of 5% from 0-100%, where location change was treated as a percentage of the maximum distance between an FA site and the membrane outline. For sensitivity of shape/size, the width and height of FA site bounding boxes was randomly altered at intervals of 5% from 0-100%, where shape/size change was considered as a percentage of the width and height of the cell, considered to be the maximum size a FA site could have. For orientation sensitivity, the angle of FA site minimum area rectangles was randomly altered at 5% intervals from 0-100%, where angle change was treated as a percentage of maximum deviation angle 90°.

For each sensitivity analysis, a plot was generated, and sensitivity values were obtained by performing a linear fit to the curves and obtaining the slope of the line. Sensitivity values were obtained for small changes between 0-10% and for large changes between 10-100%.

## Results

### 1. Cluster maps

After computing the distribution (*d*) measurement, heat maps of the FA prediction accuracy for each region of the cell showed an interpretable measure of the performance of these predictions and how FAs in some areas of the cell might be more difficult to predict than other areas. A representative example distribution heat map can be observed in Fig. 6C, where the highest accuracies can be identified in the bottom region (0.94) and middle-left region (0.93) while the upper regions had the lowest accuracy (0.44 and 0.38). Additionally, heat maps for ground truth and prediction for each cell showed a visual representation of the difference in distribution between the two images. A representative example heat maps for ground truth and prediction are shown in Fig. 6A and 6B, respectively. For these example, the bottom and middle-left regions show the most similarity between ground truth and prediction (16 vs 15 FAs for bottom region; 14 vs 13 FAs for middle-left region) while the upper regions show the largest differences (18 vs 8 FAs and 8 vs 3 FAs).

**Fig. 6:**
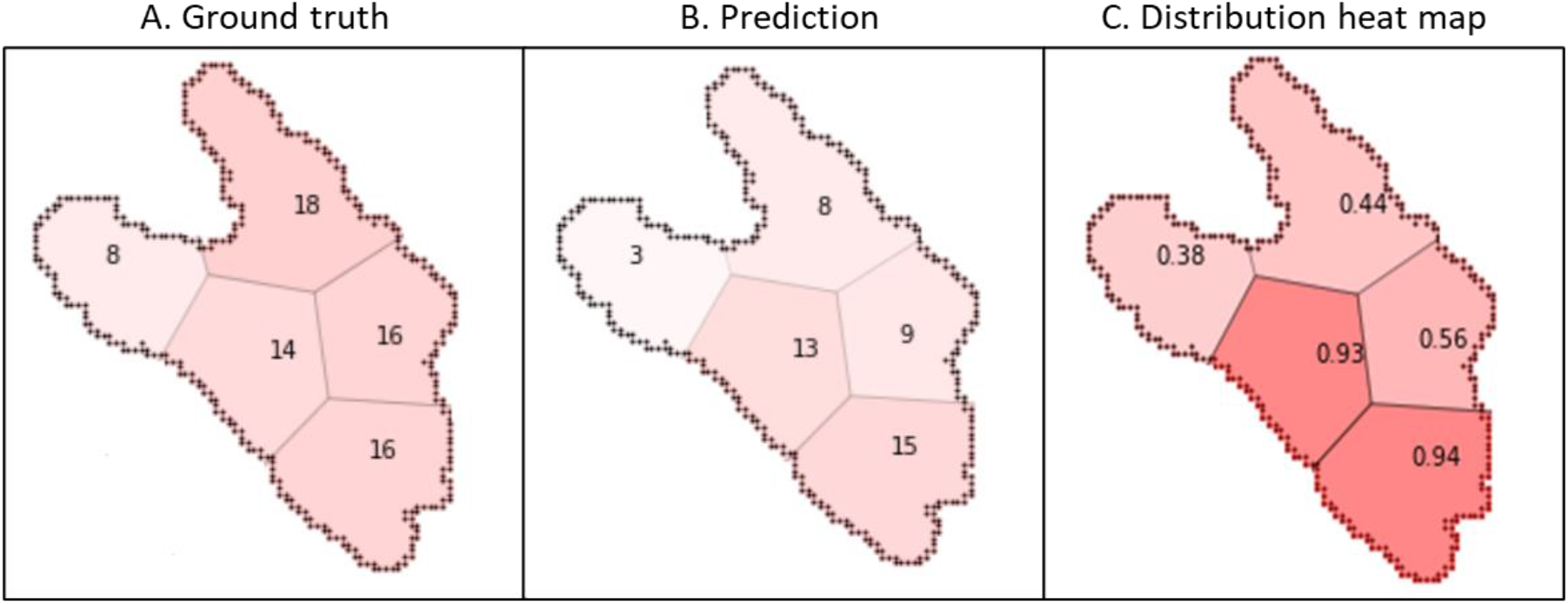
A representative sample heat map of the number of FA sites in each of the five clusters for a cell in the test set: (A) Ground truth and (B) Predicted FA sites heat map showing distribution across the five clusters where the more intense the red color, the more FAs that are present in the cluster; (C) Distribution heat map showing prediction accuracy score for each cluster where the more intense the red color, the higher accuracy.

### 2. Sensitivity analysis

Sensitivity plot for location changes is shown in Fig. 7. It can be observed that at small changes, the DPM decreases at a lower rate than all other metrics. On the other hand, at larger changes, the DPM decreases at a higher rate. Sensitivity values for this test can be seen in Table 1. DPM sensitivity was much lower (5.9E-03) than all other metrics for small changes. On the other hand, sensitivity was much higher for DPM (8.6E-03) than for all other metrics at large changes.

**Table 1.** Sensitivity values for small (0-10% dev) and large (10-100% dev) location changes in FA sites

| PCC |  | FSIM |  | IoU |  | DPM |  |
| --- | --- | --- | --- | --- | --- | --- | --- |
| 0-10%<br>dev | 10-100%<br>dev | 0-10%<br>dev | 10-100%<br>dev | 0-10%<br>dev | 10-100%<br>dev | 0-10%<br>dev | 10-100%<br>dev |
| 2.6E-02 | 4.5E-03 | 1.6E-02 | 1.6E-03 | 4.8E-02 | 1.1E-04 | 5.9E-03 | 8.6E-03 |

**Fig. 7:**
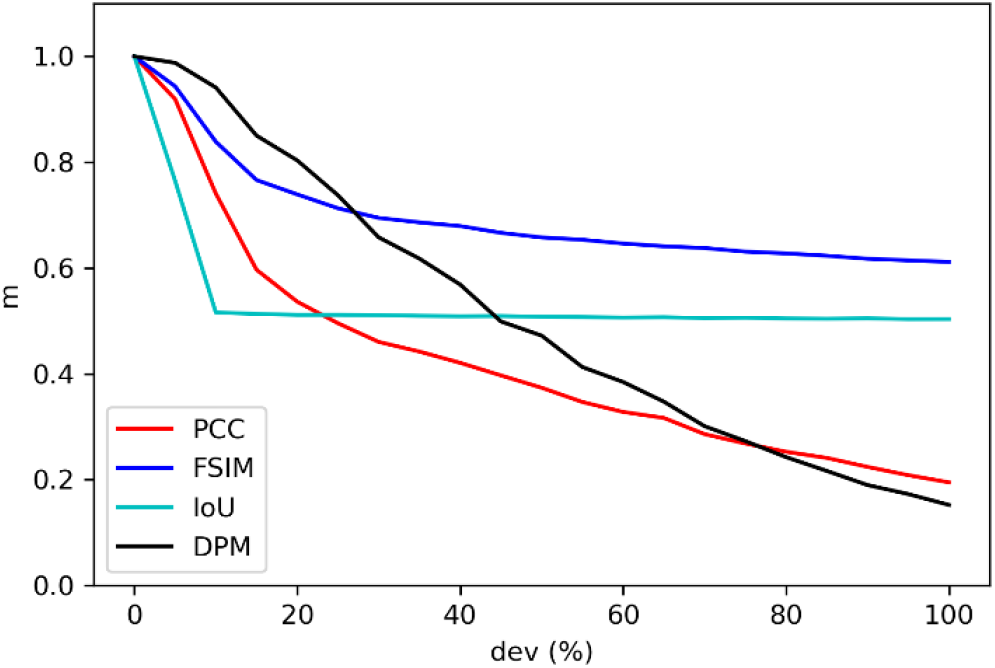
Sensitivity plot for location changes, with horizontal axis representing the deviation percentage (*dev*) and vertical axis representing metric value (*m*).

Sensitivity plot for dropping FAs is shown in Fig. 8. It can be observed that for both small and large changes, the DPM decreases at a higher and constant rate compared to all other metrics. Sensitivity values for this test can be seen in Table 2. DPM sensitivity was much higher than all other metrics and remained nearly constant for both small (9.4E-03) and large (9.9E-03) changes.

**Table 2.** Sensitivity values for small (0-10% dev) and large (10-100% dev) drops in number of FA sites

| PCC |  | FSIM |  | IoU |  | DPM |  |
| --- | --- | --- | --- | --- | --- | --- | --- |
| 0-10%<br>dev | 10-100%<br>dev | 0-10%<br>dev | 10-100%<br>dev | 0-10%<br>dev | 10-100%<br>dev | 0-10%<br>dev | 10-100%<br>dev |
| 3.3E-03 | 7.8E-03 | 3.1E-03 | 3.2E-03 | 4.7E-03 | 5.0E-03 | 9.4E-03 | 9.9E-03 |

**Fig. 8:**
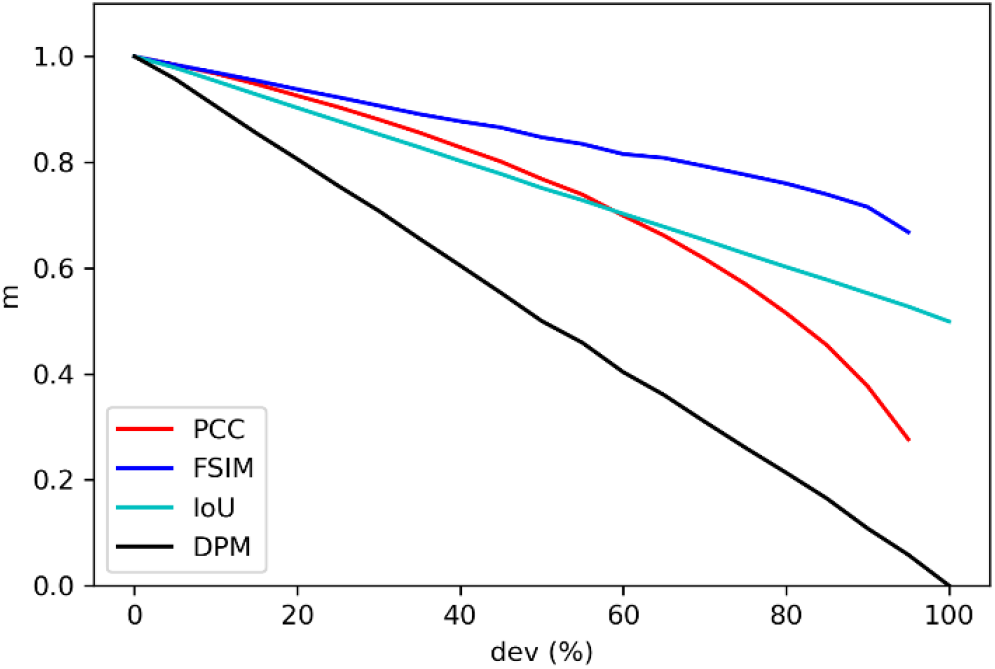
Sensitivity plot for dropping FA sites, with horizontal axis representing the deviation percentage (*dev*) and vertical axis representing metric value (*m*).

Sensitivity plot for adding FAs is shown in Fig. 9. It can be observed that for both small and large changes, the DPM decreases at a higher rate compared to all other metrics. Sensitivity values for this test can be seen in Table 3. DPM sensitivity was much higher for small (8.6E-03) than for large (4.3E-03) changes.

**Table 3.**
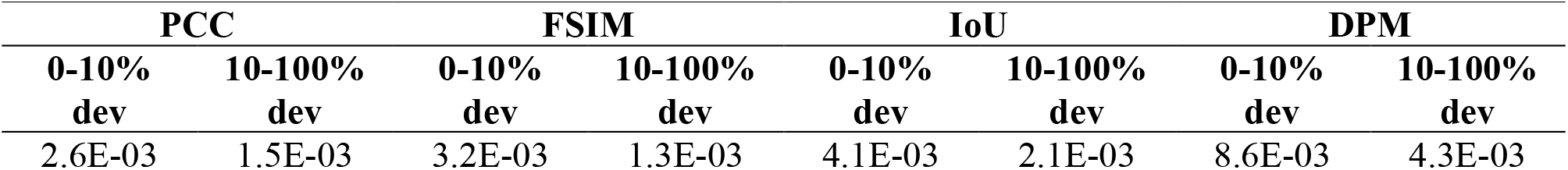
Sensitivity values for small (0-10% dev) and large (10-100%) additions in number of FA sites

**Fig. 9:**
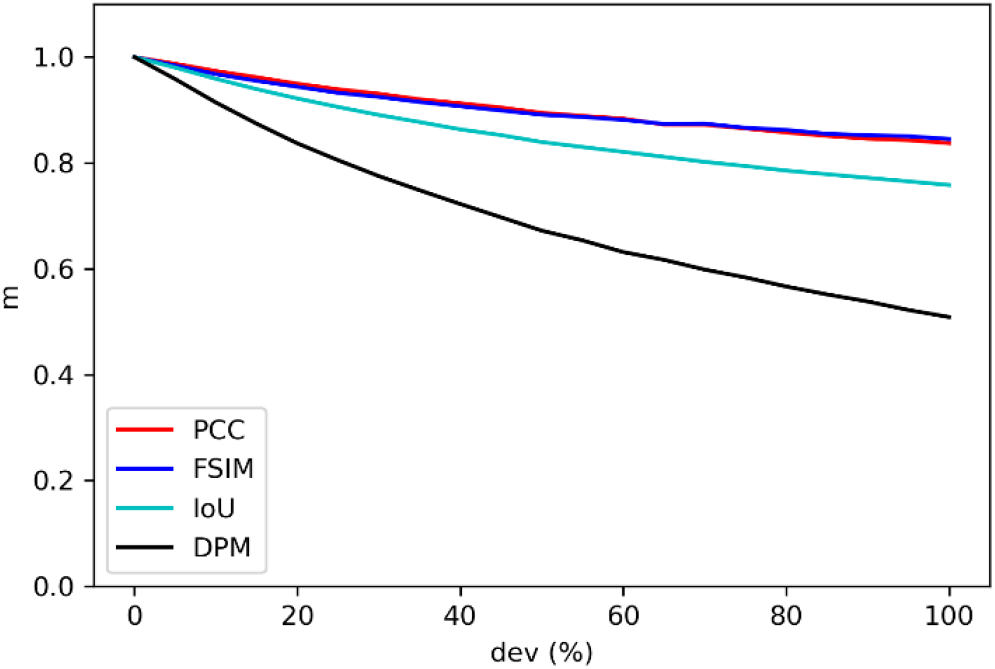
Sensitivity plot for adding FA sites, with horizontal axis representing the deviation percentage (*dev*) and vertical axis representing metric value (*m*).

Sensitivity plot for shape/size changes is shown in Fig. 10. It can be observed that for both small and large changes, the DPM decreases at a slightly higher rate compared to all other metrics. Sensitivity values for this test can be seen in Table 4. DPM sensitivity was higher than all other metrics for small (3.7E-02) and large (4.7E-03) changes.

**Table 4.** Sensitivity values for small (0-10% dev) and large (10-100% dev) shape/size changes in FA sites

| PCC |  | FSIM |  | IoU |  | DPM |  |
| --- | --- | --- | --- | --- | --- | --- | --- |
| 0-10%<br>dev | 10-100%<br>dev | 0-10%<br>dev | 10-100%<br>dev | 0-10%<br>dev | 10-100%<br>dev | 0-10%<br>dev | 10-100%<br>dev |
| 3.4E-02 | 4.1E-03 | 3.1E-02 | 1.4E-03 | 2.7E-02 | 2.9E-03 | 3.7E-02 | 4.7E-03 |

**Fig. 10:**
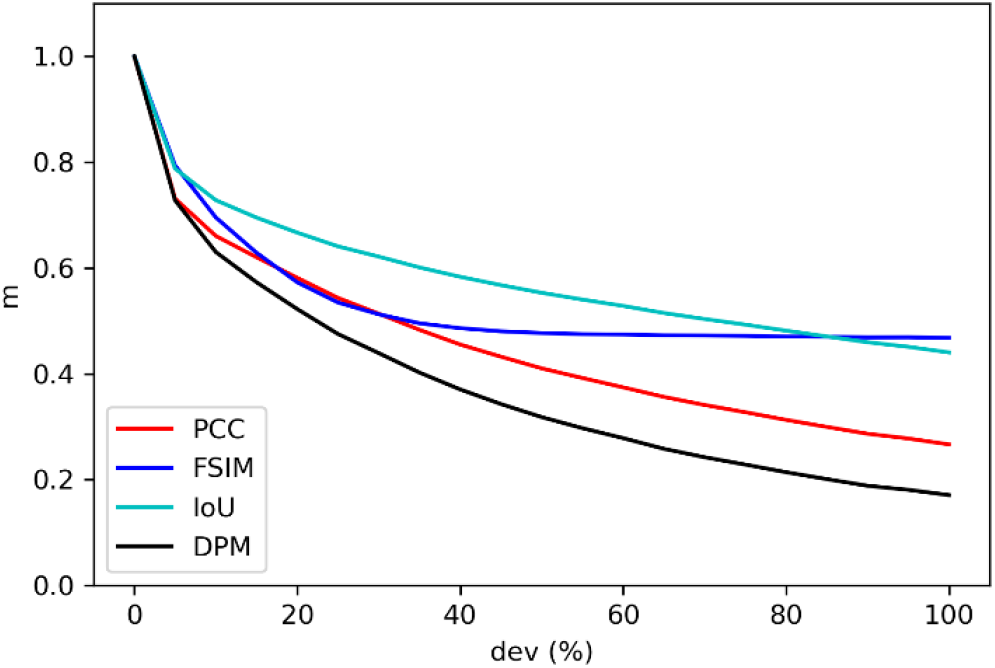
Sensitivity plot for shape/size changes, with horizontal axis representing the deviation percentage (*dev*) and vertical axis representing metric value (*m*).

Sensitivity plot for angle changes is shown in Fig. 11. It can be observed that at small changes the DPM decreases at an intermediate rate, while for large changes it decreases at a higher rate compared to all other metrics. Sensitivity values for this test can be seen in Table 5. DPM sensitivity was lower than most other metrics for small changes (9.9E-03), while it was higher than all other metrics for large changes (6.9E-03).

**Table 5.**
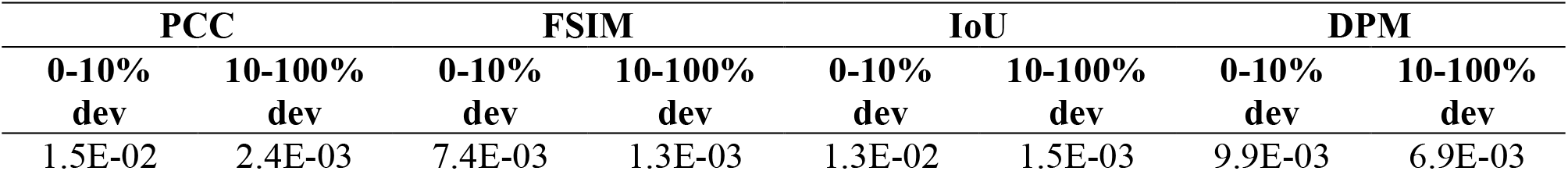
Sensitivity values for small (0-10% dev) and large (10-100% dev) angle changes in FA sites

**Fig. 11:**
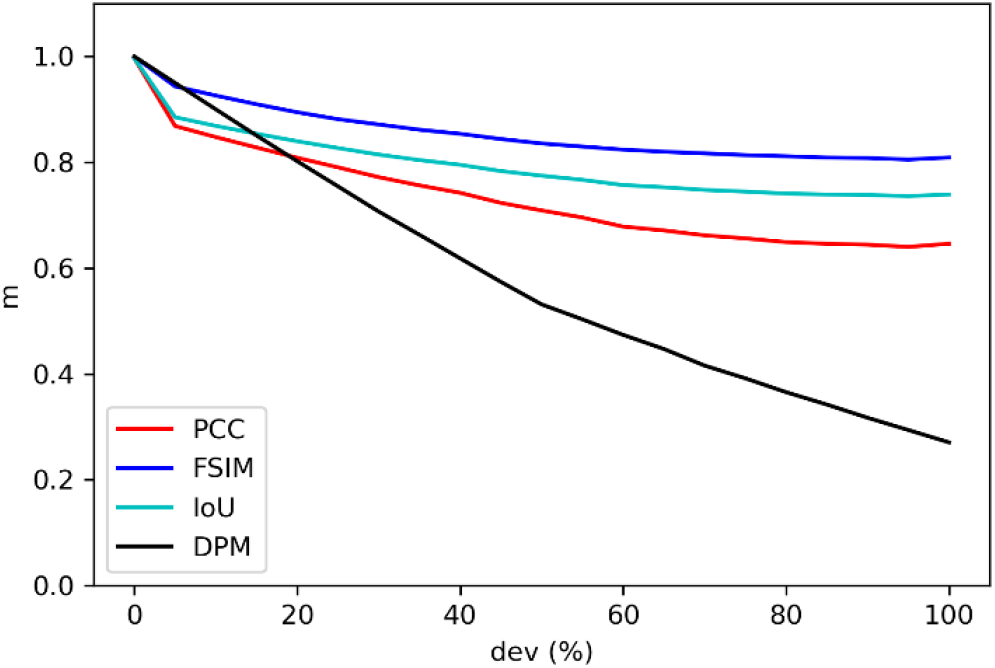
Sensitivity plot for angle changes, with horizontal axis representing the deviation percentage (dev) and vertical axis representing metric value (m).

### 3. Metric values distribution

The distribution of the values for each metric is shown in Fig. 12. FSIM and IoU showed the least variation across samples, whereas PCC and DPM showed more variation. The DPM can detect outliers in the predictions, and it also showed the longest range of values (0.36) as seen in Table 6.

**Table 6.** Mean and range values obtained across all test samples for each metric

| PCC |  | FSIM |  | IoU |  | DPM |  |
| --- | --- | --- | --- | --- | --- | --- | --- |
| Mean | Range | Mean | Range | Mean | Range | Mean | Range |
| 0.33 | 0.25 | 0.83 | 0.081 | 0.54 | 0.097 | 0.62 | 0.36 |

**Fig. 12:**
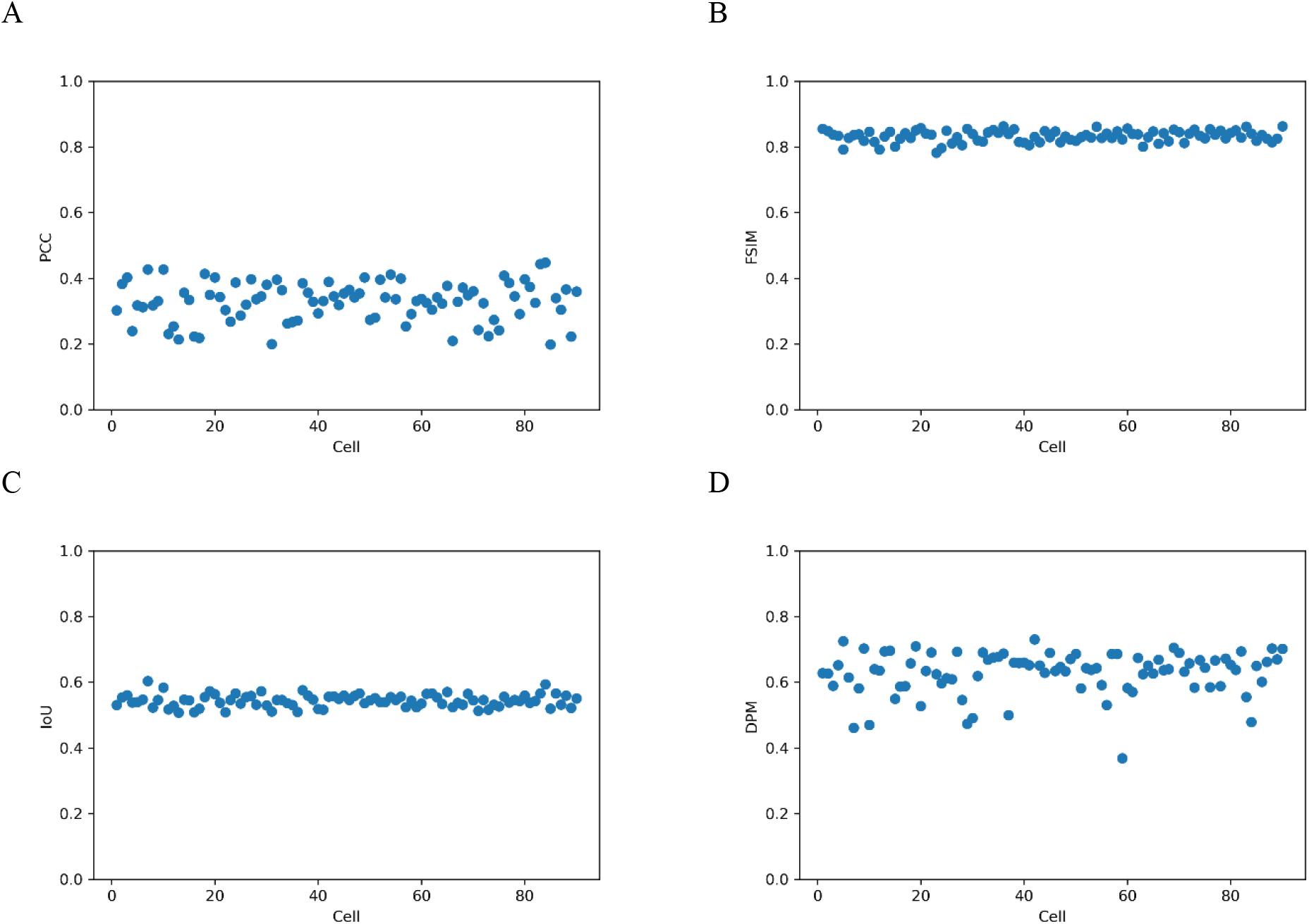
Distribution of metric values across samples in test set, where horizontal axis represents the cell number and the vertical axis the metric value for: (A) PCC; (B) FSIM; (C) IoU; or, (D) DPM.

To demonstrate the ability of the DPM to distinguish accurate predictions from inaccurate predictions, the PCC, FSIM, IoU and DPM values for two representative FA samples in the test set were compared and shown in Fig. 13. These quantitative metric values were compared to qualitative assessments for each sample. Qualitative assessment showed higher quality for Sample 2 (Fig. 13B), as more areas with roughly the same number of FA sites were identified as well as more similarities in shape/size and orientation in comparison to Sample 1 (Fig. 13A). The DPM was able to successfully distinguish that Sample 2 was quantitively a more accurate prediction than Sample 1, which is consistent with Sample 2 having a higher qualitative quality than Sample 1. On the other hand, PCC, FSIM, and IoU determined that Sample 1 was quantitively more accurate than Sample 2.

**Fig. 13:**
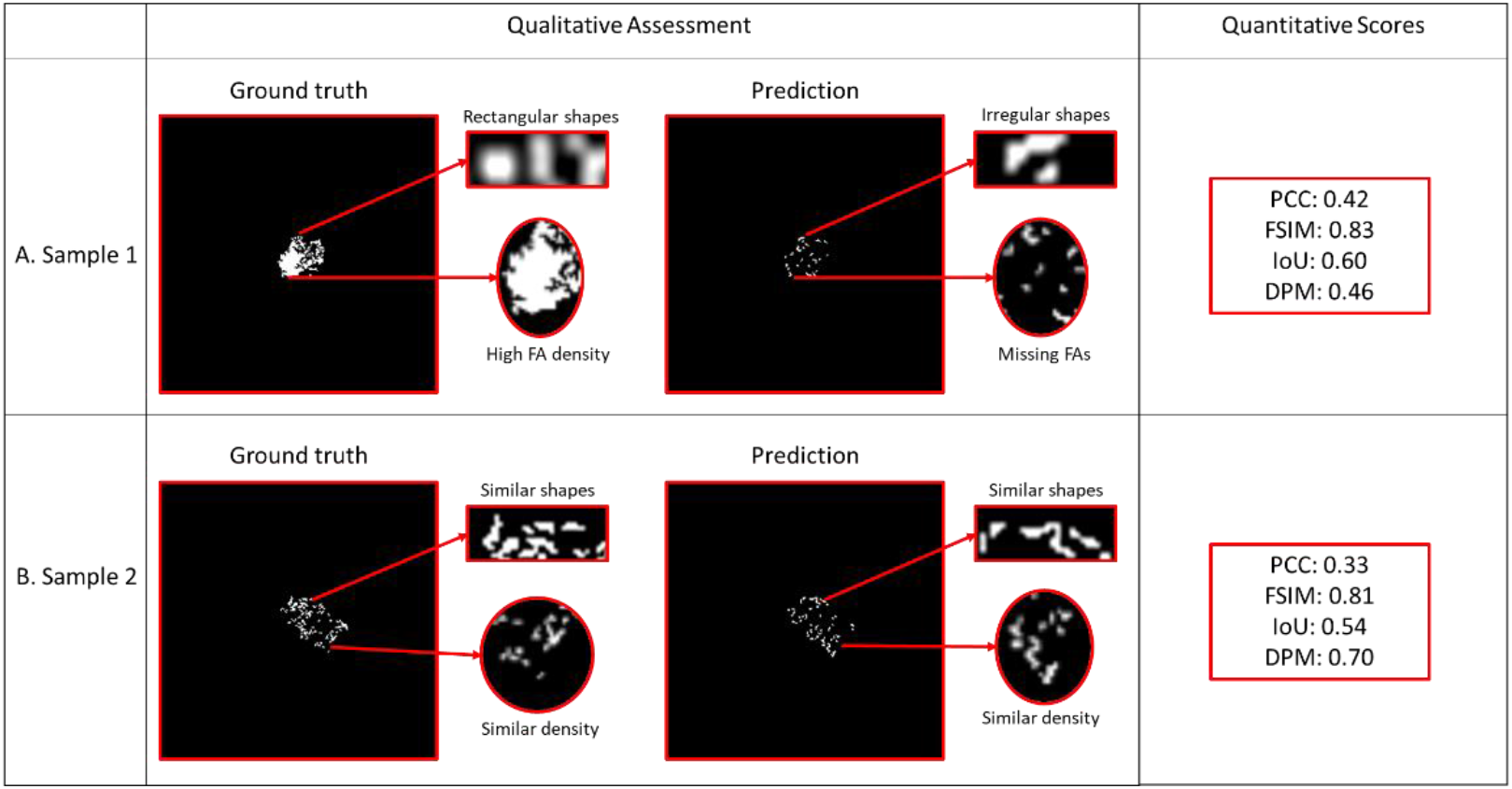
Comparison between two representative samples in test set: A. Sample 1 ground truth (left) versus prediction (right) qualitative assessment and quantitative scores; B. Sample 2 ground truth (left) versus prediction (right) qualitative assessment showing higher quality than Sample 1 and quantitative scores showing higher DPM score than Sample 1 but lower PCC, FSIM, and IoU scores.

## Discussion

The results presented show the effectiveness of DPM to calculate quantitative accuracy scores for FA site predictions that agree with qualitative assessment. Furthermore, the DPM provides a method with better sensitivity and better interpretability compared to existing metrics to determine the similarity between a predicted FA image and a ground truth image.

In terms of FA site locations, DPM has a lower sensitivity for small changes. Most of the time, predicted FAs will not be at the exact location as ground truth FAs. Therefore, it is important for the evaluation metric to have some tolerance to small differences in location. If a difference in location is small enough that an FA remains in the same region (*i*.*e*., cluster), the predicted FA location can be considered acceptable (Mullen et al., 2014). Compared to the other metrics that show drastic changes in their score for only small changes in location, the DPM allows for this tolerance. For example, PCC values drop from 1 to 0.74 for only a 10% deviation in location (Fig. 7). This drastic change can decrease the overall accuracy score, making a reasonable prediction look like a bad prediction. On the other hand, large changes in location should be clearly reflected on the value of a metric. The bigger the difference in location, the more FAs that will be located outside of the cell and, therefore, the worse the prediction. DPM has a higher sensitivity at these large deviations, whereas FSIM and IoU plateau after drastically decreasing at smaller deviations. Furthermore, for FA sites located near the edge of the cell, small deviations in location might result in the FA site being outside of the cell boundary. The DPM will tolerate small differences but will correctly penalize the prediction if the FA site is outside of the cell even if the difference in location is small.

In terms of number of FA sites, DPM had higher sensitivity to both randomly dropping and randomly adding FA sites than all other metrics that were tested. As the number of FA sites predicted can drastically affect a cell’s behavior (*e*.*g*., internal cell tension) (Mullen et al., 2014), this measurement has a high importance. While PCC, for example, shows scores as high as 0.90 after dropping 25% of FA sites, the DPM shows a score (0.75) which is more in accordance with this change in FA site number (Fig. 8). Similarly, PCC shows high scores (0.90) after increasing the number of FAs by 45%, while DPM once again shows a score (0.69) more consistent with this change in FA site number (Fig. 9). Overall, PCC, FSIM, and IoU do not change substantially for large changes in FA site number, while DPM scores better reflect these changes.

In terms of the shape/size of FA sites, the DPM had an overall slightly higher sensitivity than all other metrics that were tested. Sensitivity values were higher for small changes, consistent with the importance to detect these small changes in FA shape/size. The DPM showed lower values at 10% deviation (0.63) than PCC (0.66), FSIM (0.69), and IoU (0.72) (Fig. 10). Even more important is the ability of a metric to detect large changes in FA shape/size. The DPM showed much lower values at 80% deviation (0.21) compared to all other metrics (PCC: 0.31, FSIM: 0.47, IoU: 0.48) (Fig. 10). These measurements are important since shape/size of FAs can have a substantial impact on the overall morphology of a cell and, therefore, its behavior. For example, when looking at the mechanical behavior of a cell, the higher the substrate stiffness the larger the size of FAs and the more force they will likely produce (Haase et al., 2014).

In terms of the orientation angle of FA sites, DPM had lower sensitivity for small changes than most other metrics that were tested, while simultaneously it had higher sensitivity for large changes. This lower sensitivity is due to small differences in angle having a high impact in the other metrics tested, as these shifts in orientation have a large impact in the match of pixel values for FA sites. On the other hand, DPM shows smaller changes in its score since it directly calculates the difference between ground truth and predicted FA angle and therefore shows a more linear relationship between accuracy scores and deviation percentage. At 10% angle deviation, the DPM showed higher values (0.90) than PCC (0.84) and IoU (0.86) (Fig. 11). However, at 80% deviation all other metrics showed values much higher (PCC: 0.64, FSIM: 0.81, IoU: 0.74) than DPM (0.36) (Fig. 11). Small differences in orientation angle may not be as significant as large differences. For example, a large difference in angle can mean a drastic change in the direction of forces in a cell due to a change in the direction of flow and shear stress in the cell (Davies et al., 1994). The DPM sensitivity for angle differences is more consistent with this behavior.

A FA prediction can have deviations in location, number, shape/size, and angle with respect to its ground truth. Therefore, to evaluate the accuracy of a prediction, a metric needs to have a balanced sensitivity across all these measurements to be able of recognizing such deviations. However, it should also have the ability to capture the variation in a population. In other words, being able to discern which predictions are more or less accurate. Looking at the distribution of the metric values across test set samples (Fig. 12), the ability of DPM to capture the variation of the test samples was greater than any other metric that was tested. For instance, both FSIM and IoU show little variation in their values across samples (0.08 and 0.09 respectively), implying that all predictions were equally as good. On the other hand, PCC showed higher variation (0.24). DPM had the highest range of variation (0.36) out of all metrics that were used, showing its ability to identify a higher degree of variation within predictions leading to a better tool for validating DL training. Furthermore, the DPM provides interpretability by measuring distribution (*d*), shape/size (*s*), and angle (*a*). Across all samples in the test set, each metric showed different average values: PCC 0.32, FSIM 0.82, IoU 0.54, DPM 0.61. However, DPM provides additional information for prediction accuracy on the three specific measures (distribution, shape/size, and angle): *d* 0.61, *s* 0.57, *a* 0.67. This information can provide better insight into the strengths and weaknesses of a neural network model’s predictions. For example, results in the test set show that the neural network used has higher accuracy for predicting the distribution and angle of orientation of FAs compared to predicting the shape and size for each FA. Furthermore, Fig. 6 shows further interpretation of the distribution measurement. The heat maps provide key information into which areas of the cell have a higher prediction accuracy for FAs. For example, for some cells the network might be more accurate at predicting FAs near the center of the cell compared to the edges (as seen in the sample shown in Fig. 6).

In Fig. 13, the DPM is shown to be capable of determining which FA prediction is quantitively more accurate and consistent with qualitative assessment while other metrics do not. Sample 1 (Fig. 13A) prediction can be seen to be visually less similar to its ground truth than Sample 2 and its ground truth (Fig. 13B). However, PCC determines that Sample 1 (0.42) is a more accurate prediction than Sample 2 (0.33), FSIM determines that predictions for both samples are of similar accuracy compared to their respective ground truth images (Sample 1: 0.83, Sample 2: 0.81), and IoU also determines that Sample 1 (0.60) is a more accurate prediction than Sample 2 (0.54). On the other hand, DPM determines that Sample 2 is a more accurate prediction than Sample 1 with a score that is nearly 1.5 times different (Sample 1: 0.46, Sample 2: 0.70). This ability is crucial to the analysis of FA predictions, as these scores will be used to determine if a neural network is performing well or if further tuning is needed.

The applications of DPM are not limited to FA predictions. This metric could be extended to study the prediction or segmentation of other discrete subcellular structures. For example, mitochondrial marker *Tom20* (Johnson et al., 2017) shows a structure like that of FA sites, showing discrete structures across the cell. Prediction of such structures could be analyzed with DPM, looking at the distribution, shape/size, and angle of each mitochondrion compared to ground truth images. In addition, segmentation accuracy of small structures in CT scans could also be analyzed with this metric by comparing automated segmentation to ground truth segmentation (Ren et al., 2018). The DPM can provide better interpretability for segmentation accuracy compared to commonly used metrics such as Dice coefficient which only accounts for shape/size of structures and Hausdorff distance which only accounts for location of structures (Raudaschl et al., 2016). Furthermore, since the DPM allows the weight of each measurement (*d, s, a*) to be adjusted, the DPM allows users to tune the weight/importance of each measurement, depending on the research question of interest. For example, in the case of FA sites, mechanical behavior of a cell, such as internal cell tension, might be most affected by the number and location of FAs (Mullen et al., 2014), and, therefore, a higher weight might be given to the distribution (*d*) measurement.

## Conclusion

The proposed DPM provides a new image similarity metric that can calculate quantitative accuracy scores for DL-generated FA site predictions that are consistent with qualitative assessment. Furthermore, this method has better interpretability than the other image similarity metrics tested by measuring differences in distribution, shape/size, and angle of each FA site. The DPM can be extended to calculate accuracy of prediction and/or segmentation of other discrete structures of medical relevance.

## Code availability

Code for calculating discrete protein metric (DPM) score is on GitHub at https://github.com/mechanobiology/dpm. It includes raw focal adhesion (FA) sites ground truth and prediction samples for testing the DPM function, as well as expected DPM values for the samples and the image processing pipelines.

## Acknowledgements

Research project funded in part by the Kansas Idea Network of Biomedical Research Excellence (K-INBRE); Grant number: P20 GM103418.

## Author contributions

Conceptualization, M.C. and D.S.L.; Methodology, M.C., W.B. and D.S.L.; Software, M.C.; Data Acquisition, W.B.; Data Analysis, M.C.; Investigation, M.C.; Writing – Original Draft, M.C.; Writing – Review & Editing, M.C., W.B., and D.S.L.; Visualization, M.C.; Supervision, D.S.L.; and Funding Acquisition, M.C. and D.S.L.

## Declaration of interests

All authors declare no competing interests.

## Notes

### Competing Interest Statement

The authors have declared no competing interest.

## References

Al-Kofahi, Y., Zaltsman, A., Graves, R., Marshall, W., & Rusu, M. (2018). A deep learning-based algorithm for 2-D cell segmentation in microscopy images. BMC Bioinformatics, 19(1), 365. https://doi.org/10.1186/s12859-018-2375-z

Berginski, M. E., Vitriol, E. A., Hahn, K. M., & Gomez, S. M. (2011). High-Resolution Quantification of Focal Adhesion Spatiotemporal Dynamics in Living Cells. PLoS ONE, 6(7), e22025. https://doi.org/10.1371/journal.pone.0022025

Christiansen, E. M., Yang, S. J., Ando, D. M., Javaherian, A., Skibinski, G., Lipnick, S., Mount, E., O’Neil, A., Shah, K., Lee, A. K., Goyal, P., Fedus, W., Poplin, R., Esteva, A., Berndl, M., Rubin, L. L., Nelson, P., & Finkbeiner, S. (2018). In Silico Labeling: Predicting Fluorescent Labels in Unlabeled Images. Cell, 173(3), 792-803.e19. https://doi.org/10.1016/j.cell.2018.03.040

Davies, P. F., Robotewskyj, A., & Griem, M. L. (1994). Quantitative studies of endothelial cell adhesion. Directional remodeling of focal adhesion sites in response to flow forces. Journal of Clinical Investigation, 93(5), 2031–2038. https://doi.org/10.1172/JCI117197

Gaetani, R., Zizzi, E. A., Deriu, M. A., Morbiducci, U., Pesce, M., & Messina, E. (2020). When Stiffness Matters: Mechanosensing in Heart Development and Disease. Frontiers in Cell and Developmental Biology, 8, 334. https://doi.org/10.3389/fcell.2020.00334

Haase, K., Al-Rekabi, Z., & Pelling, A. E. (2014). Mechanical Cues Direct Focal Adhesion Dynamics. In Progress in Molecular Biology and Translational Science (Vol. 126, pp. 103–134). Elsevier. https://doi.org/10.1016/B978-0-12-394624-9.00005-1

John, J., Nair, M. S., Anil Kumar, P. R., & Wilscy, M. (2016). A novel approach for detection and delineation of cell nuclei using feature similarity index measure. Biocybernetics and Biomedical Engineering, 36(1), 76–88. https://doi.org/10.1016/j.bbe.2015.11.002

Johnson, G. R., Donovan-Maiye, R. M., & Maleckar, M. M. (2017). Generative Modeling with Conditional Autoencoders: Building an Integrated Cell. 1705.00092 [q-Bio, Stat]. http://arxiv.org/abs/1705.00092

Kärki, T., & Tojkander, S. (2021). TRPV Protein Family—From Mechanosensing to Cancer Invasion. Biomolecules, 11(7), 1019. https://doi.org/10.3390/biom11071019

Kensert, A., Harrison, P. J., & Spjuth, O. (n.d.). Transfer Learning with Deep Convolutional Neural Networks for Classifying Cellular Morphological Changes. 10.

McCarron, J. G., Lee, M. D., & Wilson, C. (2017). The Endothelium Solves Problems That Endothelial Cells Do Not Know Exist. Trends in Pharmacological Sciences, 38(4), 322–338. https://doi.org/10.1016/j.tips.2017.01.008

Mullen, C. A., Vaughan, T. J., Voisin, M. C., Brennan, M. A., Layrolle, P., & McNamara, L. M. (n.d.). Cell morphology and focal adhesion location alters internal cell stress. 12.

Nassiri, I., & McCall, M. N. (2018). Systematic exploration of cell morphological phenotypes associated with a transcriptomic query. Nucleic Acids Research, 46(19), 9.

Pedregosa, F., Varoquaux, G., Gramfort, A., Michel, V., Thirion, B., Grisel, O., Blondel, M., Prettenhofer, P., Weiss, R., Dubourg, V., Vanderplas, J., Passos, A., & Cournapeau, D. (2011). Scikit-learn: Machine Learning in Python. Journal of Machine Learning Research, 12(85), 2825–2830.

Punn, N. S., & Agarwal, S. (2020). Inception U-Net Architecture for Semantic Segmentation to Identify Nuclei in Microscopy Cell Images. ACM Transactions on Multimedia Computing, Communications, and Applications, 16(1), 1–15. https://doi.org/10.1145/3376922

Raudaschl, P. F., Zaffino, P., Sharp, G. C., Spadea, M. F., Chen, A., Dawant, B. M., Albrecht, T., Gass, T., Langguth, C., Lüthi, M., Jung, F., Knapp, O., Wesarg, S., Mannion-Haworth, R., Bowes, M., Ashman, A., Guillard, G., Brett, A., Vincent, G., … Fritscher, K. D. (2017). Evaluation of segmentation methods on head and neck CT: Auto-segmentation challenge 2015. Medical Physics, 44(5), 2020–2036. https://doi.org/10.1002/mp.12197

Ren, X., Xiang, L., Nie, D., Shao, Y., Zhang, H., Shen, D., & Wang, Q. (2018). Interleaved 3D-CNNs for joint segmentation of small-volume structures in head and neck CT images. Medical Physics, 45(5), 2063–2075. https://doi.org/10.1002/mp.12837

Rodellar, J., Alférez, S., Acevedo, A., Molina, A., & Merino, A. (2018). Image processing and machine learning in the morphological analysis of blood cells. International Journal of Laboratory Hematology, 40, 46–53. https://doi.org/10.1111/ijlh.12818

Rohban, M. H., Abbasi, H. S., Singh, S., & Carpenter, A. E. (2019). Capturing single-cell heterogeneity via data fusion improves image-based profiling. Nature Communications, 10(1), 2082. https://doi.org/10.1038/s41467-019-10154-8

Schneider, C. A., Rasband, W. S., & Eliceiri, K. W. (2012). NIH Image to ImageJ: 25 years of image analysis. Nature Methods, 9(7), 671–675. https://doi.org/10.1038/nmeth.2089

Tosh, D., & Slack, J. M. W. (2002). How cells change their phenotype. Nature Reviews Molecular Cell Biology, 3(3), 187–194. https://doi.org/10.1038/nrm761

Tschumperlin, D. J., Ligresti, G., Hilscher, M. B., & Shah, V. H. (2018). Mechanosensing and fibrosis. Journal of Clinical Investigation, 128(1), 74–84. https://doi.org/10.1172/JCI93561

Yeung, T., Georges, P. C., Flanagan, L. A., Marg, B., Ortiz, M., Funaki, M., Zahir, N., Ming, W., Weaver, V., & Janmey, P. A. (2005). Effects of substrate stiffness on cell morphology, cytoskeletal structure, and adhesion. Cell Motility and the Cytoskeleton, 60(1), 24–34. https://doi.org/10.1002/cm.20041

Yuan, H., Cai, L., Wang, Z., Hu, X., Zhang, S., & Ji, S. (2019). Computational modeling of cellular structures using conditional deep generative networks. Bioinformatics, 35(12), 2141–2149. https://doi.org/10.1093/bioinformatics/bty923

